# Molecular Basis of Mitochondrial Complex I Disruption by m.14484T>C-Induced Leber Hereditary Optic Neuropathy

**DOI:** 10.64898/2026.01.28.701874

**Authors:** Pujan Ajmera, Daniel Guion, Steven Barnes, Alfredo A. Sadun, Anastassia N. Alexandrova

## Abstract

Leber’s Hereditary Optic Neuropathy (LHON) is a rare genetic condition and severe neurological disorder characterized by dysfunctional mitochondria under extreme oxidative stress, resulting in retinal ganglion cell death and subsequent rapid bilateral loss of central vision. The m.14484T>C mutation in the ND6 subunit of mitochondrial complex I is known for inducing LHON, and is a prevalent LHON-associated mutation, yet its mechanism of impairment at the molecular level is currently unresolved. In this study, we explore the biophysical underpinnings of this mutation and its role in LHON through disruption of human complex I function. We consider, using atomistic simulations, the differential thermodynamics and kinetics of coenzyme Q10 binding between the mutant and wild-type forms, altered dynamics of the complex upon mutation, and key interactions between coenzyme Q10 and complex I binding sites. The hydrogen bond network present near and within the coenzyme Q10 binding domain, along with proper hydration of E-channel residues that couple redox chemistry to proton pumping, is found to be critical for complex I stability and quinone binding, which the ND6-centered mutation disrupts.

**Significance:** Single missense mutations to mitochondrial complex I can cause Leber Hereditary Optic Neuropathy (LHON), characterized by rapid loss of vision. However, the molecular defects produced by these mutations remain unclear, and largely inaccessible to experimental approaches. Using atomistic molecular dynamics simulations and analysis of quinone binding, we unveil the mechanism of the LHON mutation m.14484T>C in the ND6 subunit of human complex I. The mutation selectively disrupts the oxidized quinone state by increasing its distance from the terminal electron donor N2, and altering dynamic and hydration networks linked to proton pumping. This approach results in a redox-dependent model that explains experimentally observed changes in membrane potential and reactive oxygen species production upon disease-causing complex I mutations.

## Introduction

Leber’s Hereditary Optic Neuropathy (LHON) is the most common maternally inherited cause of blindness, affecting roughly 1 in 30,000 individuals worldwide.^1–3^ Typically, onset strikes teenagers or young adults, predominantly male, leading to rapid, bilateral loss of central vision.^4,5^ Clinical options are poor, often limited to idebenone, an antioxidant, or experimental gene therapies that offer genotype-dependent and incomplete rescue.^6,7^ Retinal ganglion cell axons traverse an unmyelinated segment of the retina and, lacking saltatory conduction, they rely heavily on ATP from oxidative phosphorylation, where even a modest drop in ATP production or a burst of reactive oxygen species (ROS) can precipitate apoptosis.^4,8–10^ LHON therefore represents both a clinical problem and a model for mitochondria-driven neurodegeneration.

Approximately ninety-five percent of LHON cases result from one of three point mutations in the mitochondrial genes that encode complex I subunits NADH dehydrogenase subunit 1 (ND1), subunit 4 (ND4), and subunit 6 (ND6).^11,12^ Each genotype produces a unique penetrance and recovery profile, where the ND6 mutation (m.14484T>C) is relatively mild compared to the ND4 (m.11778G>A) and ND1 mutations (m.3460G>A), with spontaneous remissions as high as 50%, while appearing more frequently in the asymptomatic population.^1,13,14^ Cybrid research has demonstrated that the ND6 m.14484T>C allele produces a drop in catalytic turnover, with measurably lowered membrane potential and elevated ROS production.^8–10,15–17^ No atomic-level mechanism linking each genotype to its unique phenotype exists.^18,19^ One of the most powerful ways to gain atomistic insight in complex I is through molecular simulations^19–21^, yet, to date only the ND1 variant has been examined with atomistic molecular dynamics simulations.^22^

Mitochondrial complex I is the largest enzyme of the electron transport chain and a critical component of aerobic energy metabolism. In humans, it is comprised of 45 subunits encompassing thousands of amino acid residues, embedded in the inner mitochondrial membrane (IMM). The catalytic cycle begins by oxidation of NADH, transferring two electrons and one proton to a flavin mononucleotide (FMN) group embedded in complex I. Electrons are then passed from FMN down eight iron-sulfur clusters which span human complex I’s peripheral arm into the Q module adjacent to the IMM. The terminal Fe_4_S_4_ cluster, N2, donates two electrons to ubiquinone (Q), and Q reduction is coupled to the uptake of two protons mediated locally, possibly by conserved His92^NDUFS2^ and Tyr141^NDUFS2^, along with two proton transfer reactions, creating ubiquinol (QH_2_). The detailed proton-routing and exact donor roles, however, remain under debate.^23–25^ The electron transfer process from NADH to ubiquinone is thermodynamically favorable due to the difference in donor-acceptor redox potential, and the free energy released is used to translocate four protons by the membrane-embedded P module into the intermembrane space, creating the proton motive force that drives ATP synthase.^26,27^ The molecular mechanism by which ubiquinone reduction is coupled to proton pumping spanning up to 200 Å remains unresolved.^20^ Prior studies have shown that this ND6 region acts as a gatekeeper for complex I hydration, potentially coupling coenzyme Q10 (CoQ10) redox chemistry to proton translocation across the membrane. In particular, redox-linked conformational shifts in ND6 are connected to opening or disruption of a continuous water-mediated proton pathway at the ND1–ND3–ND6/ND4L interface, thereby tuning when proton transfer can occur.^28,29^ m.14484T>C lies near the gate position of ND6, suggesting the mutation may perturb complex I function by altering the region that couples Q redox in the peripheral arm to proton pumping in the membrane domain.

A recent study by Rigobello and colleagues modeled the m.14484T>C mutation for the first time in molecular dynamics through coarse-grained simulations of a truncated complex I structure, absent of the CoQ10-binding region.^30^ They uncovered that the m.14484T>C mutation, which is a single methionine-to-valine mutation on the TM3 helix of the ND6 subunit (M64V^ND6^), leads to stiffening of dynamics in the E-channel responsible for linking CoQ10 redox to proton translocation. What remains unresolved is how Q/QH_2_ interact with the mutation site and how the mutation might perturb Q/QH_2_ binding. Multiple studies have shown that motions in the CoQ10 site lead to changes in the E-channel^31,32^, but atomistic simulations of the m.14484T>C mutation in a complex I model with the CoQ10 site present have yet to be explored with different redox states of Q. Here, we show that m.14484T>C disrupts a critical hydrogen bond network between ND6 and NDUFS2, altering the binding orientation of Q and disfavoring productive reduction of ubiquinone. The mutation destabilizes complex I dynamics and hydration in a redox state-dependent manner that is critical for connecting Q redox to the proton pumping mechanism.

## Results & Discussion

### Sequence conservation analysis of the ND6 mutation

Evolutionary constraints reflect the influence of selection pressures that restrict a protein’s mutational space, i.e. which sites can tolerate substitution and to what degree. To explore such constraints possibly present in complex I ND6 and contextualize the importance of the 64^ND6^ site for ND6 function, we performed multiple sequence alignment (MSA) of ND6 mammalian protein orthologs obtained from OrthoDB and assessed the MSA for positional variability (**Methods**). The conservation profile (**Figure 1C**) revealed discrete peaks of high conservation (positional variability score closer to 1) interspersed with valleys of low conservation (score closer to 0), regions in the ND6 protein. To concretely identify the most conserved region of ND6, we further defined the most conserved segment as the contiguous region that maximizes total excess conservation relative to the alignment-wide average. This threshold-free definition avoids imposing an arbitrary conservation cutoff or minimum length, while still permitting occasional local drops in conservation within an otherwise constrained region.^33,34^ Using this approach, we identified a segment spanning amino acid positions 15–85 with a mean conservation substantially higher than the global average (0.84 vs. 0.69). This region is enriched in hydrophobic residues (**Figure S1**), consistent with its proposed role as a hydration gate whose conformation depends on the redox state of Q. The LHON-associated ND6 variant m.14484T>C is located within this most conserved portion of the protein, at position 64 in both the MSA and human ND6 sequence. Overall, the analysis indicates that selection pressure is highest across residues 15–85 of MT-ND6, placing the LHON mutation site near the center of the most functionally constrained segment of the protein.

**Figure 1:**
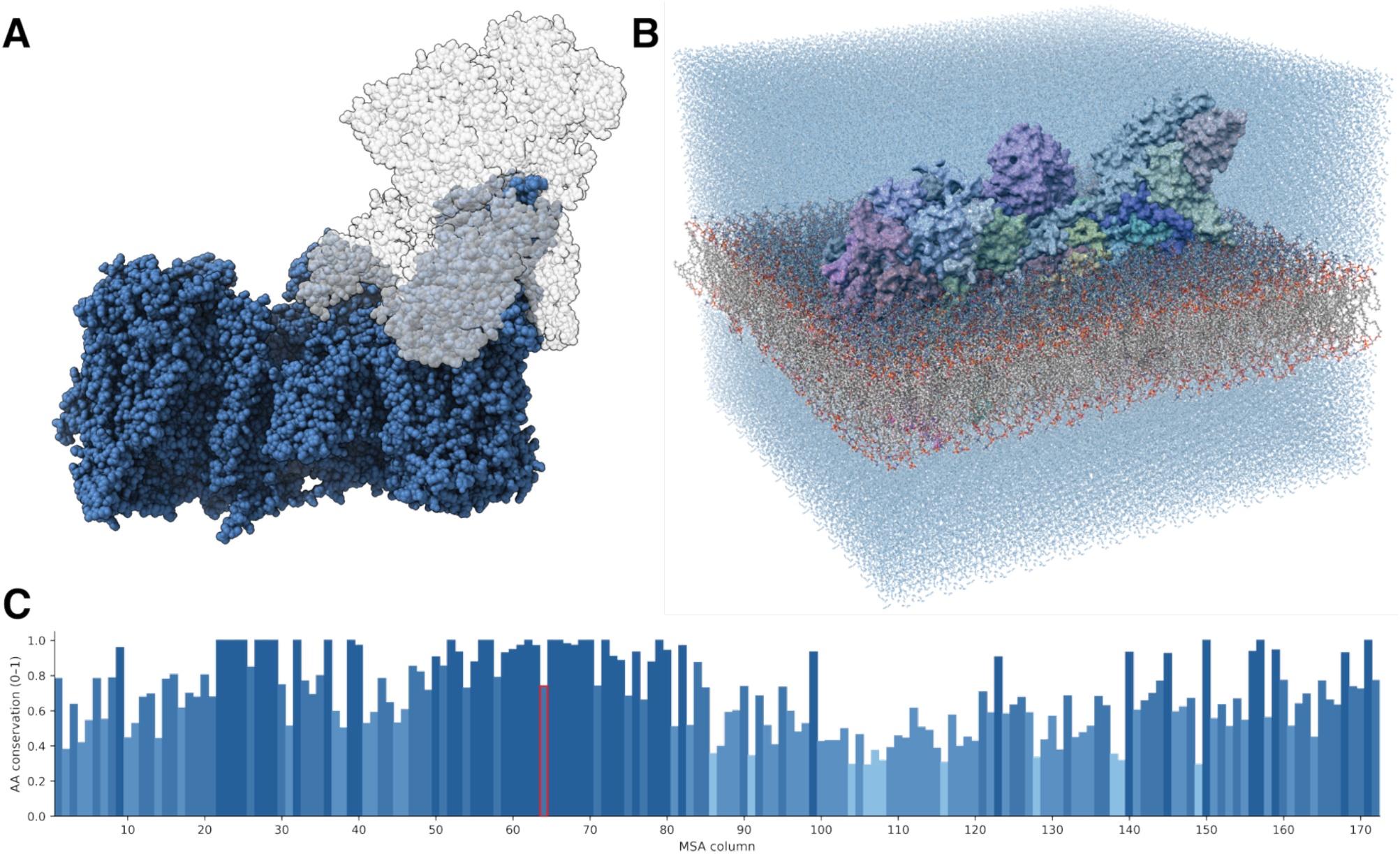
(A) The full human complex I from the cryo-EM structure 5XTD, with the chains used in the simulations shown in color. (B) The periodic image of the complex I model used in this study, including membrane and Fe/S cluster components (C) Conservation of ND6 amino acid identity across mammals (based on mammalian ortholog dataset from OrthoDB) (**Methods**). Each alignment column is represented as a vertical bar whose height corresponds to Sj (0–1), with bar color mapped to a five-level blue color scale from low to high conservation and the disease-associated site in MT-ND6 outlined in red.

### Thermodynamics and kinetics of Q/QH2 binding using non-equilibrium simulations

To assess how the m.14484T>C mutation perturbs quinone binding energetics in complex I, we performed alchemical free energy perturbation (FEP) calculations comparing WT and M64V^ND6^ for Q and QH_2_. For FEP, M64V^ND6^ was introduced through an alchemical transformation (**Methods**) using the thermodynamic cycle shown in **Figure 2A** for both unbound and bound Complex I, using previously equilibrated structures (**Methods**). Solving for the thermodynamic cycle, we computed the difference in binding free energies between WT and M64V^ND6^ Complex I as follows:

**Figure 2:**
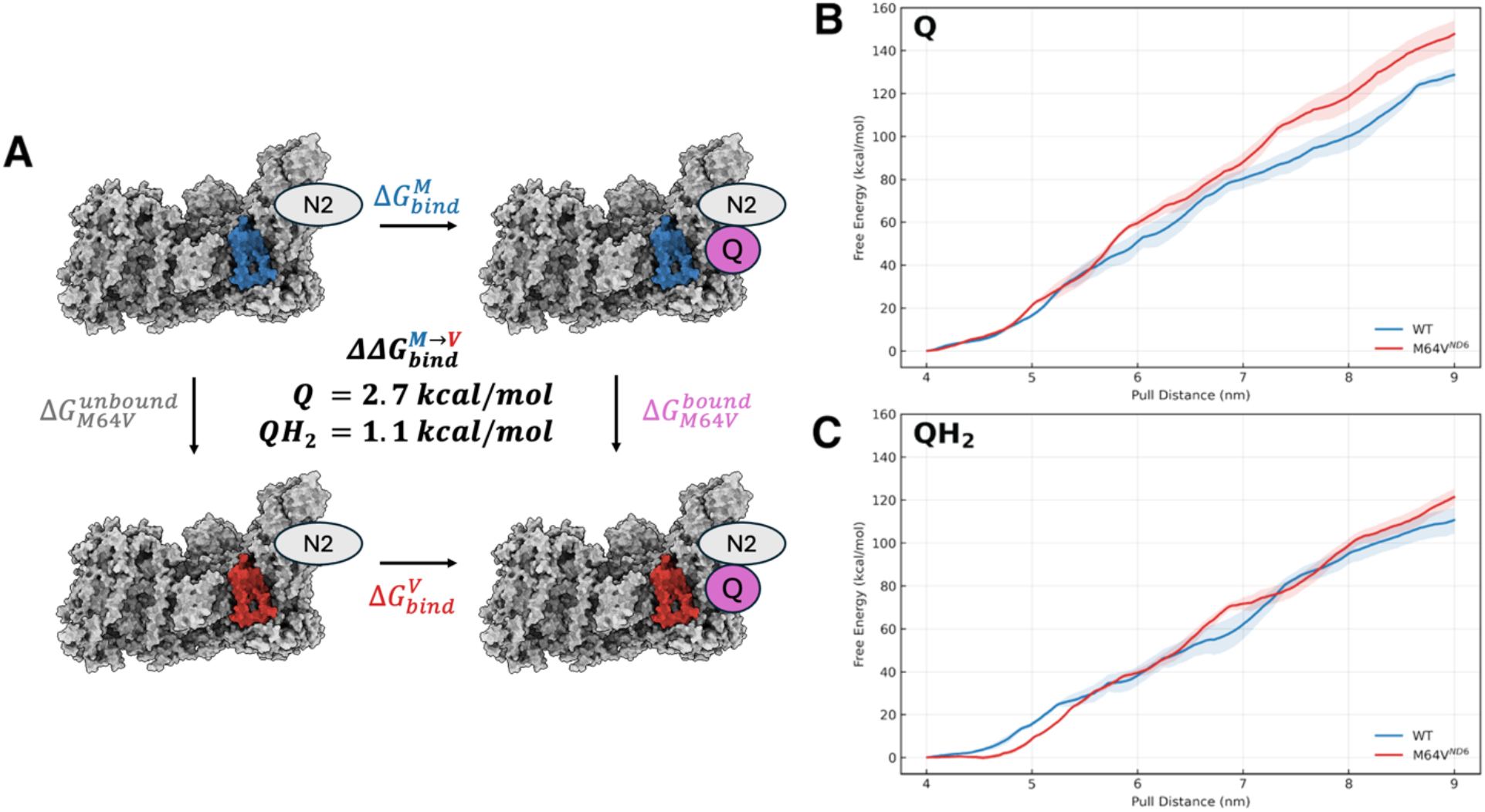
(A) Thermodynamic cycle employed to compute the relative binding free energy (ΔΔG) between wild-type (WT, blue) and M64V (red) mutant systems for Q and QH_2_ using free energy perturbation (FEP) simulations. (B) Jarzynski-averaged work profile for the egress of Q, derived from steered molecular dynamics (SMD) simulations (C) Jarzynski-averaged work profile for the egress of QH_2_, also obtained via SMD simulations.

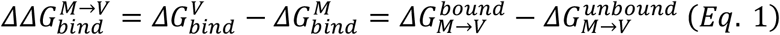

For the oxidized quinone, Q binds less favorably to the mutant Complex I than to WT, with *ΔΔG*^*M*→*V*^_*bind*_= 2.7 kcal/mol. By contrast, for QH_2_, the corresponding value is 1.1 kcal/mol, which lies within the typical uncertainty of relative binding free energy calculations (often ∼1–2 kcal/mol under favorable conditions).^35,36^ As such, we do not interpret the QH_2_ result as evidence for a substantial difference in binding free energy. Taken together, these results suggest a redox-state-specific effect of the mutation, with M64V^ND6^ selectively destabilizing the Q-bound state relative to WT by 2.7 kcal/mol.

Separately, steered molecular dynamics (SMD) simulations were performed using a moving harmonic restraint on the distance between the N2 Fe_4_S_4_ cluster and CoQ10 tail, pulled from 40Å to 90Å. Assessment of the kinetic barrier for Q and QH_2_ egress was performed using SMD, and the required free energy for pulling the CoQ10 is described by the Jarzynski equality:

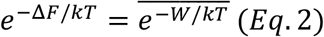

In wild-type Complex I, the SMD simulations reveal that less energy is required to remove QH_2_ from the binding cavity compared to Q, following the logic of Complex I’s catalytic cycle (**Figures 2B,2C**). Upon mutation, however, Q egress is impaired, with Q motion in M64V^ND6^ accumulating significantly more energetic penalty compared to WT (**Figure 2A**), while QH_2_ egress appears less impacted by the mutation (**Figure 2B**). To relate the pulling coordinate to headgroup motion, we mapped the SMD trajectories onto the distance between the CoQ10 headgroup and the N2 cluster (**Figure S2**). Across all four systems, the tail group moved substantially during the first 10–12 Å of pulling while the headgroup remained near its initial binding-site position. In the mutant Q-bound system, the headgroup began from a larger N2 separation and moved closer to N2 after ∼10 Å of tail pulling, consistent with a more deeply buried initial pose than in the other three systems.

Taken together, the FEP and SMD simulations indicate that M64V^ND6^ perturbs the quinone-binding landscape in a redox-dependent manner. The clearest effect is observed for the oxidized quinone, which is both thermodynamically destabilized and kinetically hindered from leaving the mutant than WT complex I. By contrast, differences in QH_2_ are comparatively modest within the states and pulling pathway examined here. Collectively, these results suggest that the mutation disrupts the balance between substrate binding and turnover by favoring a nonproductive Q pose with hindered egress, which may impede efficient catalytic cycling in complex I. In comparison with prior atomistic simulations of the LHON-associated ND1 mutation^22^, Q in M64V^ND6^ likewise appears to occupy a perturbed binding landscape. However, unlike the ND1 mutant, M64V^ND6^ does not produce the same degree of quinol kinetic trapping during egress, suggesting that the dominant defect in m.14484T>C is not impaired QH2 release.

### Equilibrium modeling of ND6 mutation through dynamic cross-correlation and hydration gate analysis

To probe the dynamic perturbations to the complex that occur upon CoQ10 binding, and how introducing mutations alters those dynamic motions, we performed long equilibrium molecular dynamics simulations of complex I with QH_2_, Q, and unbound quinone for both WT and M64V^ND6^ forms. Stability of the simulations were verified using RMSD (**Figure S3**). Dynamic cross correlation (DCC) matrices, which determine the degree of dynamic relationship between different protein chains, were computed for 200ns trajectories (**Methods**). Strongly correlated motions (DCC ∼ +1) indicate comparatively rigid domains, whereas strongly anticorrelated motions (DCC ∼ -1) indicate hinge motions or open/closing breathing modes of the complex. Introduction of M64V^ND6^ to complex I results in a dramatic loss of correlated motions in the complex when Q is bound to the cavity shown in **Figure 3A** and **Figure 3B**. Most of the disruption is a loss of positive correlation rather than a gain in anticorrelation, implying dynamic instability. The loss in correlation observed occurs at a long-range, with smaller changes near the Q-binding NDUFS2 domain. Upon reduction to ubiquinol, the mutation to complex I has a near-reversed effect on correlated motions in the protein (**Figure 3C, D**), with an increase in long-range correlated motions between chains near from NDUFS2 and those further away in M64V^ND6^ as observed in WT. Short-range correlations near the binding site are only somewhat disrupted, connecting to the lack thermodynamic and kinetic penalties that quinone binding incurs upon mutation. This mutation therefore has a redox-dependent effect on the dynamics of complex I, where correlated motions presumably required for proper Q binding and motion to the second binding site post reduction to QH_2_ in WT are missed in M64V^ND6^. Upon quinone binding regardless of the N2 redox state, complex I is found to increase in dynamic cross correlation (**Figure S4 & S5**) in the wild-type complex, indicating collective domain movement when Q is bound relative to the free protein. Ligand-protein binding has been previously found to impact protein dynamics through changes in cross-correlation^37,38^, and prior studies have demonstrated the impact of mutations on protein dynamic correlations and loss of enzyme function.^39,40^

**Figure 3:**
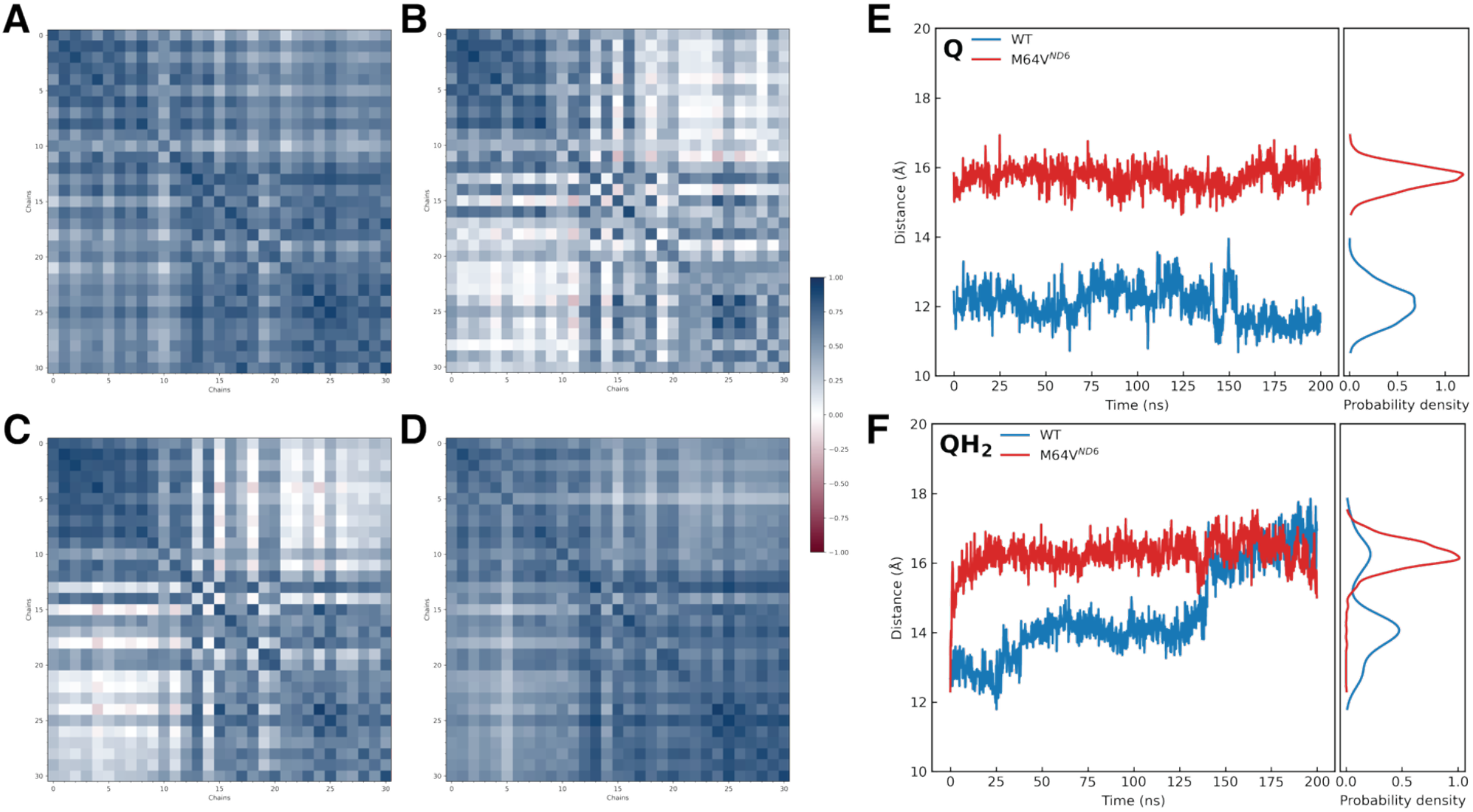
Dynamic cross correlation (DCC) matrices for (A)WT bound to Q, (B) M64V^ND6^ bound to Q, (C) WT bound to QH_2_, and (D) M64V^ND6^ bound to QH_2_. DCC values in red indicate anticorrelation, and blue indicates correlation. Values are group averaged by chain, and DCC matrices are sorted by distance to Q-binding site. Quinone head to N2 distances for (E) Q-bound and (F) QH_2_-bound forms over the simulation, with corresponding kernel densities.

We also quantified the equilibrium separation between N2 and the Q headgroup in both WT and the M64V^ND6^ mutant. In WT complex I with Q bound, the Q-N2 distance is 12.1 ± 0.5 Å (**Figure 3E**), consistent with prior reports that the canonical reactive pose places Q 10–14 Å from N2, known as binding site 1, to enable efficient electron transfer between redox pairs.^31,41^ Without reaching binding site 1, Q cannot undergo redox with N2 to initiate the redox-driven proton pump. In contrast, we observe a Q-N2 distribution shifts to larger distances in M64V^ND6^ and centered at 15.7 ± 0.3 Å, indicating that Q is substantially less likely to achieve the < 14 Å tunneling distance typically required for Q reduction (**Figure 3E**).^42^ Upon reduction, QH_2_ is expected to relax away from binding site 1 position into distal parts of the Q chamber.^20,31^ Consistent with this, we find that QH_2_ resides at a mean separation of 14.6 ± 1.3 Å in WT, whereas QH_2_ in M64V^ND6^ remains close to where Q sits in the mutant binding site, at 16.2 ± 0.5 Å (**Figure 3F**). Upon inspection of the Q and QH_2_ binding poses in WT, we observe that Q’s binding pose is consistent with binding site 1 placement while QH_2_’s binding pose at the end of the 200ns WT MDs is consistent with presence at binding site 1’, where ubiquinol is expected to position itself post-reduction, farther away from N2 and Tyr141^NDUFS2^ to prevent reverse electron transfer^31^. For both Q and QH_2_ in M64V^ND6^, however, the CoQ10 occupies a site buried within NDUFS2 that is not present in WT, where the loop harboring His92^NDUFS2^ is more disordered in M64V^ND6^ compared to WT, and distances between CoQ10 and Tyr141^NDUFS2^ do not increase upon reduction (**Figure S6**). Taken together, these results are consistent with the previous non-equilibrium modeling of Q and QH_2_ binding and, in concert with the DCCM results, indicate that Q-dependent complex I dynamics are disrupted while QH_2_-dependent dynamics are more preserved with the M64V^ND6^ mutation compared to WT. The ∼4 Å change in distance would result in a c.a. 250-fold decrease in electron transfer rate upon mutation (see **Supplementary Notes**). Prominently, these results indicate that the mutant is comparatively incompetent in reducing Q to QH2. In these conditions, the mutant will overproduce ROS to the surplus of electrons unused in the reduction.

A complete picture of the M64V^ND6^ mode of complex I dysfunction requires understanding of the atomic-level perturbations that the mutation induces in the protein. From a hydrogen-bond occupancy analysis, we observe that M64V^ND6^ disrupts a hydrogen bond that normally stabilizes the TM3 α-helix in WT. We observe this disruption propagate as traceable structural changes from subunit ND6 into NDUFS2, where the Q headgroup binds, and we assign this change as the underlying biophysical perturbation responsible for altered Q binding and dynamics (**Figure 4)**. Specifically, at the mutation site, we observe slight breaking of the interaction between the backbone carbonyl oxygen of residue 64^ND6^ (O) and the backbone amide of G68^ND6^ (HN) (hydrogen bond occupancy in MD found as 0.77 versus 0.52 in WT-Q and M64V^ND6^-Q, respectively). We see loss of this interaction perturbs the positioning of TM3^ND6^ and increases conformational heterogeneity in the TM3–TM4^ND6^ connecting loop (residues 75–85) (**Figure 4A;** see **Figure S7** and **Figure S8** for root-mean squared deviation of full ND6 subunit and TM3– TM4^ND6^ connecting loop, respectively, in WT-Q, WT-QH_2_, M64V^ND6^-Q, M64V^ND6^-QH_2_). In the WT, this loop forms an inter-subunit salt-bridge with NDUFS2 through E77^ND6^ and K78^NDUFS2^ (**Figure 4B**), which is lost in M64V^ND6^ (**Figure 4C**) (hydrogen bond occupancy in MD found as 0.54 versus 0.03 in WT-Q and M64V^ND6^-Q, respectively). Comparable residue pairs at the ND6– NDUFS2 interface have been identified across several species and are shown to control a confirmational switching network that links Q redox activity to proton translocation across the IMM.^21^ In NDUFS2, His92^NDUFS2^ is a canonical hydrogen-bonding partner to Q^23–25^, which lies on a β-hairpin whose turn contains a short α-helical insert in the WT **(Figure 4D)**. In the mutant, this element collapses into a fully disordered loop **(Figure 4D)**. As a result, His92^NDUFS2^ pulls Q deeper into complex I and away from N2, shifting Q further into the NDUFS2 pocket **(Figure 4D)**. Correspondingly, there is a marked reduction in hydration throughout the E-channel (**Methods**) of the mutant in the Q-bound state compared to the wild type, whereas no such difference is observed between the mutant and WT in the QH_2_-bound state (**Figure 4E**; residue breakdown for E-channel hydration provided in **Figure S9**).

**Figure 4.**
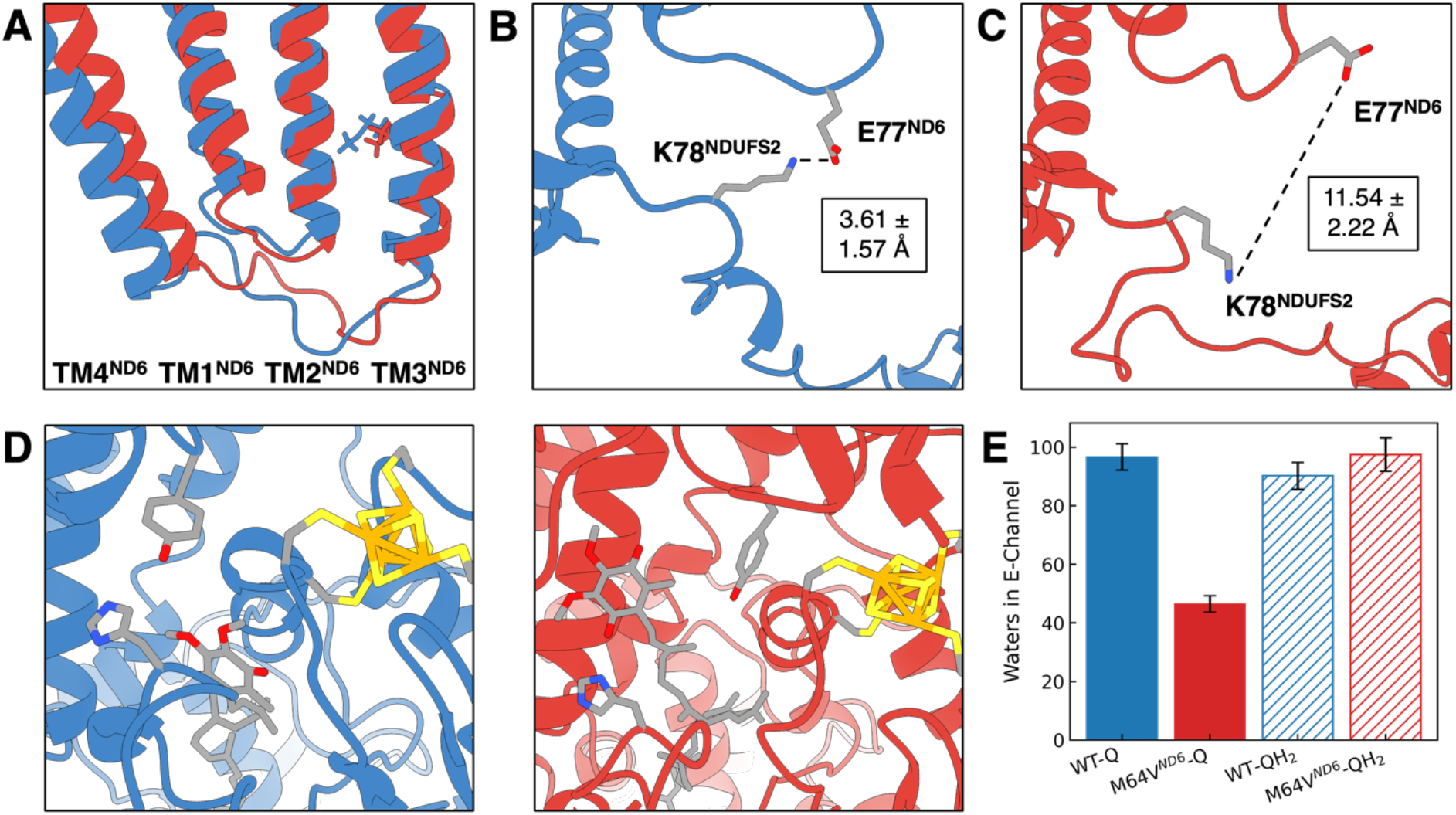
Structural consequences of the M64V^ND6^ substitution and its impact on Q-site remodeling. (A) Loss of the stabilizing *i*→*i*+4 hydrogen bond between 64^ND6^ O and 68^ND6^ HN disrupts TM3^ND6^ packing and increases conformational heterogeneity in the TM3–TM4^ND6^ loop. (B) In WT-Q, this loop forms an intersubunit salt bridge with NDUFS2 through E77^ND6^ and K78^NDUFS2^, with (C) contacts that are lost in the M64V^ND6^-Q. (D) Perturbation of the ND6–NDUFS2 interface collapses the β-hairpin/α-helical insert harboring His92^NDUFS2^ into a disordered loop, shifting Q deeper into the NDUFS2 pocket and away from N2. (E) This reorganization reduces E-channel hydration selectively in the Q-bound state.

The decreased hydration observed in Q-bound state of the mutant relative to WT arises primarily from a loss of waters surrounding E-channel residues in the ND1 and NDUFS2 subunits, particularly Glu202^ND1^, Glu227^ND1^, and His92^NDUFS2^, which are consistently less hydrated in M64V^ND6^-Q than in WT-Q (**Figure S9**). In contrast, we detect little to no comparable difference in hydration between M64V^ND6^-QH_2_ and WT-QH_2_ **(Figure S9)**. Prior cryo-EM studies of mammalian complex I reveal that Q-bound complex I demonstrates markedly increased water-mediated Glu202^ND1^–Glu227^ND1^ interactions, whereas, in the apo state, Glu202^ND1^ adopts an alternative rotamer that disrupts this interaction. The Glu202^ND1^–Glu227^ND1^ pair has previous been implicated in a Grotthuss-like proton-transfer process in which CoQ10 redox chemistry modulates local hydration near the Q site, thereby gating CoQ10-redox–triggered proton transfer into the ND1 E-channel and onward toward the membrane domain’s proton-pumping machinery.^29,43,44^ These observations suggest that M64V^ND6^ weakens or disrupts this water-mediated coupling between the Q site and the ND1 E-channel in the mutant in complex I and provide mechanistic rationale for impaired proton-coupled function.

## Conclusion

In the present study, we examined how the LHON-causing m.14484T>C mutation alters quinone binding, positioning, and coupled dynamics in human mitochondrial complex I. We observed across simulations the impact of the M64V^ND6^ mutation to be redox-state dependent, with the largest differences between mutant and WT observed in the Q-bound state, whereas the QH_2_-bound state showed comparatively smaller perturbations. We found that the M64V^ND6^ mutation destabilizes Q binding<u>binging</u> near N2 and impairs Q egress from complex I. Conversely, the kinetics and thermodynamics of QH_2_ binding are similar between mutant and WT, suggesting that binding energetics of the Q-bound state are a dominant factor of the mutant’s phenotype. Similarly, Q occupied a position farther from N2 in M64V^ND6^ than in WT, which is a shift expected to reduce the efficiency of productive electron transfer and thereby disfavor Q reduction relative to WT. Consistent with these findings, long-range correlated dynamic motions and hydration of the E-channel are lost in a redox-dependent manner where solely Q bound to M64V^ND6^ results in drastic changes in dynamics and decreased hydration relative to WT. These results were not present in the QH_2_-bound state.

We find the mutation site itself slightly destabilizes the TM3 helix of ND6, which serves as a hydration gate linking Q-site redox in the peripheral arm to conformational changes in the membrane domain that drive proton pumping. This localized destabilization propagates from subunit ND6 to NDUFS2 as M64V^ND6^ disrupts a key intersubunit salt bridge, altering NDUFS2 loop dynamics and disrupting Q-headgroup positioning, causing it to shift deeper into the Q-binding cavity and away from N2. Additionally, hydration around the proton-transfer pair Glu202^ND1^–Glu227^ND1^, which is proposed to couple Q-site redox to the proton-relay into the membrane domain, is disrupted by M64V^ND6^, specifically in the Q-bound state but not in the QH_2_-bound state. Overall, these findings indicate that the m.14484T>C mutation disrupts the normal redox-coupled dynamics of complex I, altering the Q-dependent coupled redox-proton pumping mechanism, which would reduce the generation of the proton motive force essential for mitochondrial and other cellular functions.

Compared with atomistic simulations of the more penetrant LHON-associated ND1 mutation m.3460G>A, our results suggest that M64V^ND6^ (m.14484T>C) does not impair ubiquinol egress from the complex I Q cavity to the same extent. Although m.14484T>C alters the Q site, its dominant effect appears to be on elements of the coupling machinery that link Q reduction to proton pumping, thereby indirectly perturbing the Q chamber and disproportionately affecting the Q-bound state. These findings are consistent with a coupling-deficient phenotype for m.14484T>C, rather than the stronger kinetic trapping and greater ROS production observed in computational and experimental m.3460G>A models.^8,9,17,22^ Importantly, M64V^ND6^ increased the separation between the Q headgroup and N2, which is expected to reduce productive electron transfer, but did not produce the extent of Q-chamber remodeling previously reported for ND1 m.3460G>A. Together, these simulations suggest that ND6 m.14484T>C perturbs complex I through a mechanism distinct from the more severe Q-site remodeling observed for ND1 m.3460G>A. This difference may help explain why m.14484T>C is often associated with a milder bioenergetic phenotype and lower clinical penetrance.

## Methods

We constructed the computational models used in this study using the cryo-EM model of intact human complex I (PDBID 5XTD), removing the distal half of the peripheral arm, including segments upstream of the N6a/N6b/N2 clusters to avoid computationally prohibitive calculations, while retaining tunnels responsible for proton transfer and the binding of CoQ10. The resulting POPC/POPE/cardiolipin (50:35:15) bilayer, 150 mM KCl solvent, and protein sum to ∼1.1 million atoms, constructed with CHARMM-GUI^45^. Coenzyme Q10 was docked in the oxidized form (Q) in wild-type form with a reduced N2 cluster (WT-rN2-Q). Protonation states were assigned using Propka^46,47^ at pH 7 to accurately model membrane protein charges^48^. Parameters from the CHARMM36 force field^49^ were used for all atoms besides the Fe_4_S_4_ clusters, for which parameters were obtained from previous studies.^50,51^

To relax contacts from the cryo-EM model, minimizations were performed on WT-rN2-Q with restraints: first minimizing just the lipids, then including waters and ions, then minimizing the full system, using OpenMM^52^. Then, heating and equilibration were performed over 10ns to bring the system to 310K. After equilibration, the M64V^ND6^ mutation was introduced *in silico* using CHARMM, and QH_2_ replaced Q for both the WT and M64V^ND6^ constructs, yielding a total of four systems: WT-rN2-Q, WT-oN2-QH_2_, M64V^ND6^-rN2-Q, and M64V^ND6^-oN2-QH_2_, where oN2 indicates the corresponding oxidized state of N2. We performed an additional 10ns of equilibration for all four systems to respond to changes in oxidation states and mutation. All equilibrations were done in the NPT ensemble. From these structures, the WT-rN2, WT-oN2, M64V^ND6^-rN2, and M64V^ND6^-oN2 constructs were prepared by removing Q/QH_2_ from the final structures of the previous equilibration, and additionally equilibrating for 10ns at 310K. Final structures from this and the previous equilibration were used for further analysis.

The equilibrated Q/QH_2_ bound structures were used to begin force pulling simulations. The pulling coordinate used, to have a consistent reference, was the distance between the N2 Fe_4_S_4_ cluster and the tail carbon of Q/QH_2_. The distance was first equilibrated at 40Å for 10ns for the WT and M64V^ND6^ variant. A harmonic force (*k*=8 kcal/mol/Å^2^) was used to pull the quinone along the tunnel axis at 0.5 Å/ns for 50 Å, yielding 100ns of simulation for each force pulling run. Samples every 1ns of the fixed distance equilibration were used to seed force pulling, providing 10 replicates for each WT-rN2-Q, WT-oN2-QH_2_, M64V^ND6^-rN2-Q, and M64V^ND6^-oN2-QH_2_, using OpenMM for simulating the steered molecular dynamics. Work profiles were converted to free energies via the Jarzynski equality^53^. Standard errors were computed via bootstrapping with replacement^54^, where random trajectories were averaged, and the corresponding standard deviations were used to obtain error bars.

Free energy calculations were performed using GPU-accelerated FEP with CHARMM in NAMD 3.0.1^55^, using the dual-topology protocol. The alchemical perturbation corresponded to the protein mutation, with the goal of assessing the binding free energy of complex I to either Q or QH_2_ upon mutation M64V^ND6^ (m.14484T>C). Because reliable FEP requires adequate pre-equilibration before introducing the alchemical perturbation^36^, dual-topology systems were constructed from the final frames of 200 ns equilibrium MD simulations of WT-rN2-Q, WT-rN2, and WT-oN2 (*vide infra*). In these systems, the methionine and valine side chains were both retained at residue 64, with hybrid parameters taken from top_all36_hybrid.inp in the VMD topology library. Mutant structures were generated with VMD Mutator 1.5 and psfgen 2.0^56^. A special procedure was required for the WT-oN2-QH_2_ system. During equilibrium MD, QH_2_ occupied an unstable position farther from N2 and showed a tendency to move toward the Q-chamber entrance, consistent with product egress after reduction. By contrast, in M64V^ND6^-oN2-QH_2_, QH_2_ remained in a trapped pose similar to that observed in M64V^ND6^-rN2-Q. Because FEP is difficult to converge when the ligand samples substantially different positions, we tested whether WT-oN2-QH_2_ could also support this trapped pose. To do so, we back-mutated the 200 ns endpoint structure of M64V^ND6^-oN2-QH_2_ to WT and performed an additional 200 ns equilibration. In this back-mutated WT construct, QH_2_ remained stably trapped within the Q-chamber, and this equilibrated endpoint was used to build the WT-oN2-QH_2_ dual-topology system. Accordingly, the resulting ΔΔ*G*_*bind,M*64*V*−*WT*_ for QH_2_ should be interpreted narrowly as the mutation-dependent change in binding free energy for this specific trapped QH_2_ pose, rather than for the full ensemble of WT QH_2_ configurations. This treatment was preferred over attempting FEP from a highly positionally heterogeneous WT-oN2-QH_2_ state, which would likely suffer from poor phase-space overlap and inadequate convergence. Production FEP simulations were carried out using 80 alchemical windows with a cosine-eased schedule (see **Supplementary Notes**). In each window, 500 ps of equilibration was followed by 2 ns of data collection, with energies recorded every 500 fs. Free energy differences were estimated with Bennett acceptance ratio using a zeroth-order Gram-Charlier expansion as implemented in VMD ParseFEP^57^. Convergence plots for all 80 windows are provided in **Figure S10-13**.

Previously equilibrated structures for the quinone-bound forms (WT-rN2-Q, WT-oN2-QH_2_, M64V^ND6^-rN2-Q, M64V^ND6^-oN2-QH_2_) were subjected to 200ns of production simulations each, using OpenMM. The structures without quinone bound (WT-rN2, WT-oN2, M64V^ND6^-rN2, and M64V^ND6^-oN2) were subjected each to 100ns of production simulations, observing well-converged RMSD within that timescale (**Figure S4**). Further simulation and analysis details can be found in the **Supplementary Notes**.

## Acknowledgements

D.G. acknowledges support from the CQSE Graduate Fellowship from the Center for Quantum Science and Engineering at the University of California, Los Angeles, supported by NSF Grant 2125924. S.B. and A.A.S. acknowledge support from an unrestricted grant from Research to Prevent Blindness, Inc. to the UCLA Department of Ophthalmology. S.B. acknowledges support from NIH R01 EY033905 and Glaucoma Research Foundation Shaffer Grant Award. A.A.S. acknowledges support from the International Foundation for Optic Nerve Diseases (IFOND). Computational resources were provided through the NSF ACCESS Grant BIO240118, for all simulations performed in this work. Molecular graphics and analyses were performed with UCSF ChimeraX, developed by the Resource for Biocomputing, Visualization, and Informatics at the University of California, San Francisco, with support from National Institutes of Health R01-GM129325 and the Office of Cyber Infrastructure and Computational Biology, National Institute of Allergy and Infectious Diseases.

