## Supporting Information for "Molecular Basis of Mitochondrial Complex I Disruption by m.14484T>C-Induced Leber Hereditary Optic Neuropathy"

Table of Contents

| | |
|---|---|
| Supplementary Notes – Methodology and Analysis | S1-S2 |
| Supplementary Notes – Marcus-Hush Theory and Redox Cycle | S3 |
| Figure S1 | S4 |
| Figure S2 | S5 |
| Figure S3 | S6 |
| Figure S4 | S7 |
| Figure S5 | S8 |
| Figure S6 | S9 |
| Figure S7 | S10 |
| Figure S8 | S11 |
| Figure S9 | S12-S13 |
| Figure S10 | S14 |
| Figure S11 | S15 |
| Figure S12 | S16 |
| Figure S13 | S17 |
| References | S18 |

**Supplementary Notes – Methods and Analysis**

*Simulation settings for equilibrium trajectories, and force-pulling in OpenMM*

All simulations in OpenMM were performed in the NPT ensemble at 310K and 1.01325 bar. A Langevin thermostat was used to maintain temperature, with the available Monte Carlo barostat for constant pressure. A 2 fs integration timestep was used. Non-bonded interactions were treated with a 10 Å cutoff. Rigid bonds for hydrogens were enforced using the SETTLE algorithm. For treatment of long-range electrostatics, the Particle Mesh Ewald approach was used with a 5E-4 error tolerance to set the grid.

*Simulation settings and analysis details for Free Energy Perturbation (FEP) in NAMD*

To ensure proper sampling along FEP, we employed a cosine-eased λ-schedule to define the intermediate states between λ = 0 and λ = 1. In this scheme, Windows are generated by taking uniform steps in an angle x from 0 to π and mapping them to λ with a half-cosine transform: $\lambda = (1 - \cos x)/2$. This produces a smooth, symmetric schedule with $\lambda = 0$ at $x = 0$, $\lambda = 0.5$ at $x = \pi/2$, and $\lambda = 1$ at $x = \pi$. Because the slope $d\lambda/dx = (1/2)sin(x)$ is near zero at the endpoints and largest near the middle, the resulting λ-increments are small (dense windows) near $\lambda = 0$ and $\lambda = 1$ and largest (sparser windows) near $\lambda = 0.5$, concentrating sampling where endpoint behavior is typically most sensitive. FEP simulations were performed in the NPT ensemble at 310 K and 1.01325 bar. Temperature was maintained using a Langevin thermostat, and pressure was controlled using a Nosé–Hoover Langevin piston barostat. A 2 fs integration timestep was used. Non-bonded interactions were treated with a 12 Å cutoff, and a 10 Å switching function was applied to smooth the cutoff boundary for electrostatic and van der Waals interactions. Rigid bonds for hydrogens were enforced using the SETTLE algorithm. For treatment of long-range electrostatics, the Particle Mesh Ewald approach was used with a 1 Å grid spacing.

*Production simulation analysis*

The dynamic cross-correlation matrix (DCCM) of the residues was computed via the MD-TASK package[1], with modifications made for parallelization given the system size. Averages of the DCCM values across and between protein chains provided the final DCCM maps shown in this work, to simplify the data from correlations between c.a. 5000 residues to between 31 chains. The final frame from each equilibrium MD trajectory was used for the structural analysis in **Figure 4A-D**. Trajectory-based of RMSD, distances, hydrogen bond occupancy, and hydration were carried out using the last 50ns of the 200ns equilibrium trajectories (full 200ns simulation for DCCM and Q/$QH_2$-N2 distances) and analyzed using the MDAnalysis library in Python.[2] E-channel hydration (**Figure 4E**) was evaluated by counting unique water molecules within 5 Å of key residues defined by Kim et al.[3] including ND4L (Glu34, Glu70), ND6 (Met64/Val64), ND3 (Asp66), ND1 (Glu143, Glu192, Glu202, Glu206, Glu227), and NDUFS2 (His92, Tyr141, Asp193, Arg111, Arg115).

*Bioinformatics analysis*

Protein orthologs of mammalian MT-ND6 were obtained (N = 124) from OrthoDB[4] and aligned using MAFFT v7 with default parameters.[5] Entries sharing the same OrthoDB organism taxid were collapsed to a single representative, selecting the least gapped sequence. The alignment was filtered with trimAl (v1.4) to remove sequences with insufficient coverage of the shared aligned region, where we required that a sequence overlap the well-aligned core by at least 80 amino-acid positions. "Well-aligned" positions were defined as columns in which at least 75% of sequences contained a residue rather than a gap (using the -resoverlap 0.75 -seqoverlap 80 criteria). Gap-rich columns were further trimmed using trimAl's gappyout procedure.[6] The final alignment (N = 101), including variation in residue identity and type was visualized (**Figure S1)** using the ggmsa package.[7] Position-based Henikoff sequence weighting was then applied to reduce the influence of redundant or closely related sequences on conservation estimates.[8] For each alignment column, the number of distinct amino acid types and their counts were used to assign fractional contributions to each sequence. These per-column contributions were then summed across the alignment to yield a total weight for each sequence. The weights were normalized so that their sum equaled the number of sequences. Weighted amino acid frequencies were then computed for each column using the 20 standard proteogenic amino acids. Gaps were excluded from the normalization. A small pseudocount ($10^{-3}$) was added to avoid zero probabilities.[9] The final curated alignment was then assessed for positional variability using Shannon entropy[10] computed from the weighted frequencies $p_{j,a}$ of amino acid $a$ at column $j$:

$$H_j = -\sum_a p_{j,a} \log_2 p_{j,a} \ (Eq. S1)$$

To place entropy on a 0–1 conservation scale independent of the number of amino acid states, we defined a per-position conservation score such that a perfectly conserved column has $S_j = 1$ and a maximally diverse column that is uniform over 20 amino acids has $S_j \sim 0$:

$$S_j = 1 - \frac{H_j}{\log_2(20)} \ (Eq. S2)$$

The most conserved stretch of residues in a threshold-free manner was defined as the contiguous set of positions [a, b] that maximizes the sum $\sum_{j=a}^{b} x_j$, where $x_j$ is the excess conservation at position j relative to the average ($\bar{S}$):

$$x_j = S_j - \bar{S} \ (Eq. S3)$$

The maximizing segment was identified using a linear-time maximum-subarray algorithm[11,12] implemented in Python.

**Supplementary Notes – Marcus-Hush Theory and Redox Cycle**

We can utilize the following Marcus-Hush equation[13], with the approximation introduced by Dutton[14] for estimating the electron transfer rate as a function of the redox potential $\Delta G_{ET}$ and the distance between redox centers R, with the approximate reorganization energy $\lambda = 0.7 eV$[14]:

$$log_{10}(k_{ET}) = 15 - 0.6R - 3.1 \frac{(\Delta G_{ET} + \lambda)^2}{\lambda} \ (Eq. S4)$$

We are careful here to evaluate R as the edge-edge distance, and now the ratio of the log of electron transfer rates between WT and M64V$^{ND6}$ (the latter simply referred to as "MUT") as:

$$log_{10}\left(\frac{k_{ET,WT}}{k_{ET,MUT}}\right) = -0.6(R_{WT} - R_{MUT}) - \frac{3.1}{\lambda}\left[\left(\Delta G_{ET,WT} + \lambda\right)^2 - \left(\Delta G_{ET,MUT} + \lambda\right)^2\right]$$
$$= -0.6(R_{WT} - R_{MUT}) - \frac{3.1}{\lambda}\left(\Delta G_{ET,WT} - \Delta G_{ET,MUT}\right)\left(\Delta G_{ET,WT} + \Delta G_{ET,MUT} + 2\lambda\right)$$

The thermodynamic cycle used in the main text can also be similarly used to obtain a redox cycle, where, if we assume the free energy of reduction of the ubiquinone in WT and $M64V^{ND6}$ can be well-mapped to the electron transfer driving force, gives a negligible $\Delta G_{ET,WT} - \Delta G_{ET,MUT}$ of ~1 kcal/mol (0.043eV). From **Figure 3**, $R_{WT} - R_{MUT}$ for the Q-bound state is roughly 4 Å, and therefore we approximate the ratio of electron transfer rates to be solely dependent on the change in distance:

$$log_{10}\left(\frac{k_{ET,WT}}{k_{ET,MUT}}\right) \approx 2.4$$

This results in a c.a. 250-fold decrease in electron transfer rate.

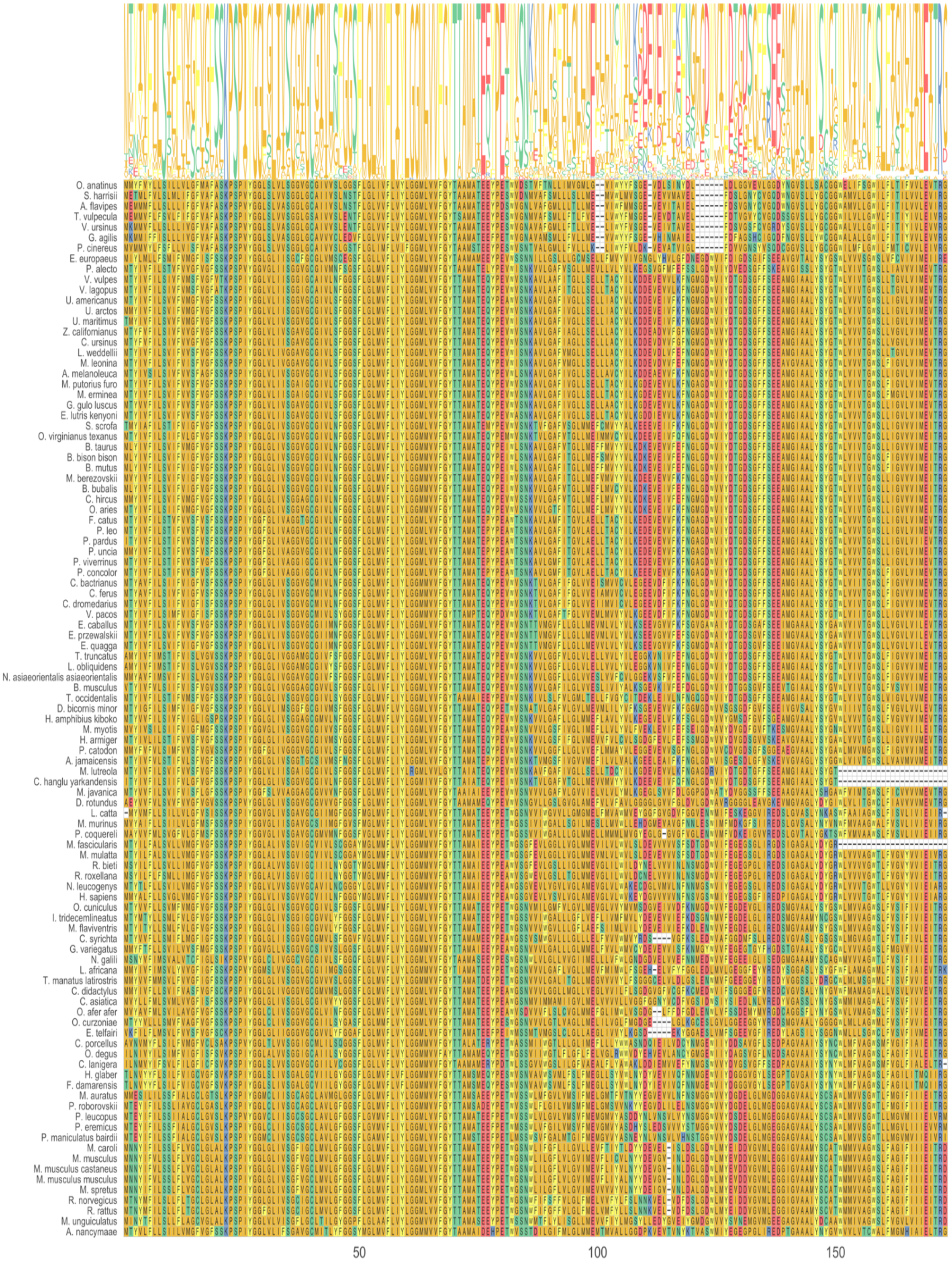

**Figure S1**: MSA of Mammalian ND6 Orthologs obtained from OrthoDB (Public Orthologous Group ID: 420823at40674 / level taxid: 40674) (**See Methods**)

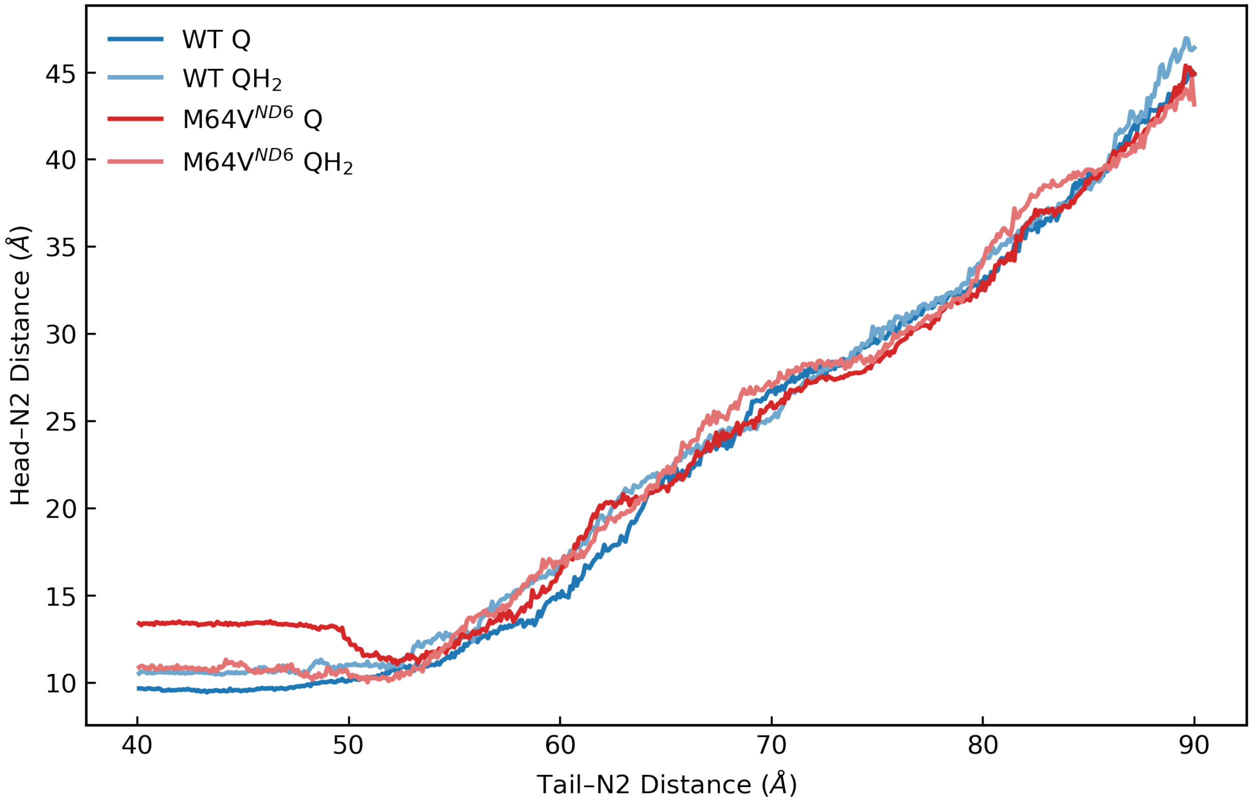


**Figure S2**: Head versus tail distances to N2 during SMD molecular dynamics.

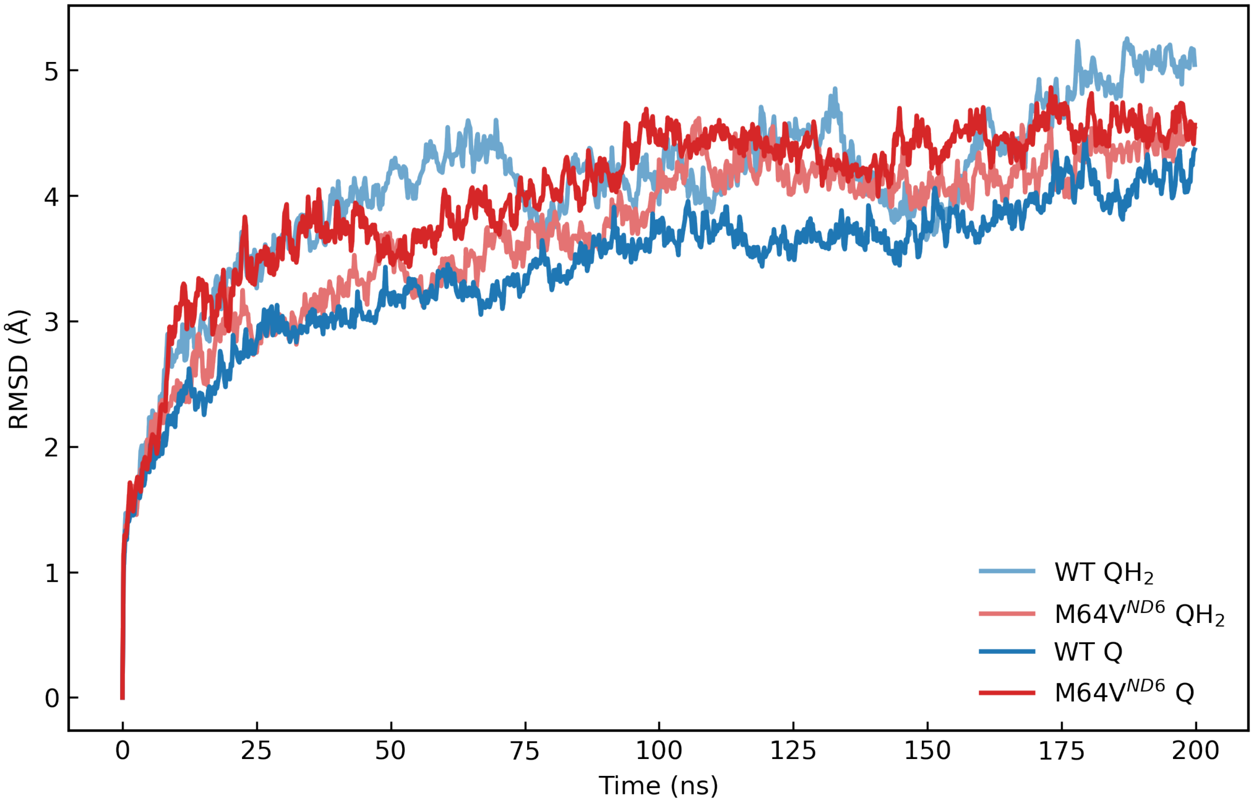


**Figure S3**: Backbone root-mean-square deviation (RMSD) of the whole protein complex, relative to the initial frame, for each equilibrium molecular dynamics trajectory in the Q- and $QH_2$-bound states.

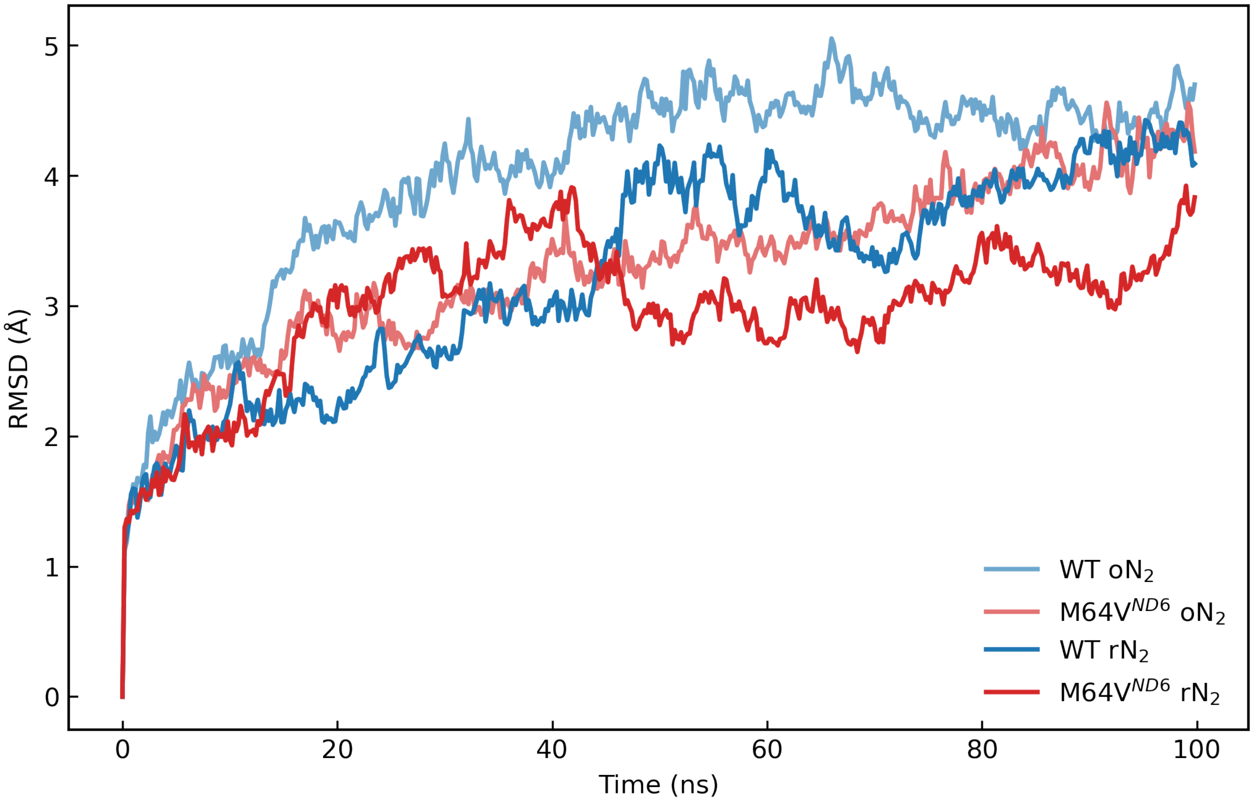


**Figure S4**: Backbone root-mean-square deviation (RMSD) of the whole protein complex, relative to the initial frame, for each equilibrium molecular dynamics trajectory in the ligand-free states containing reduced N2 (rN2) or oxidized N2 (oN2).

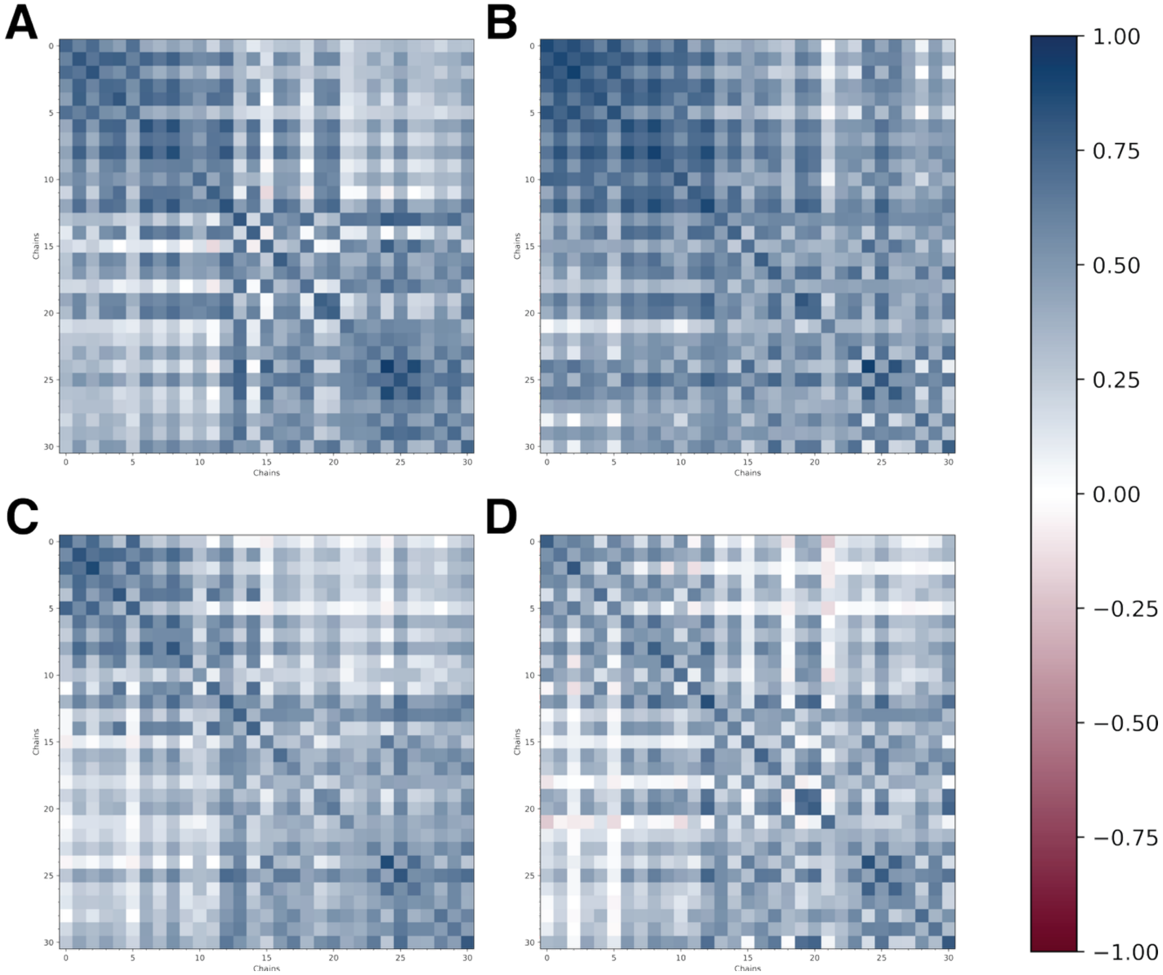


**Figure S5:** DCC matrices for unbound Q-forms of Complex I for (A) WT with reduced N2, (B) M64V$^{ND6}$ with reduced N2, (C) WT with oxidized N2, and (D) M64V$^{ND6}$ with oxidized N2. Positive values (blue) indicate stronger correlated motions, and negative values (red) indicate stronger anticorrelated motions. Chain numbers in axes are grouped as described in **Methods** and **Figure 3** in the main text.

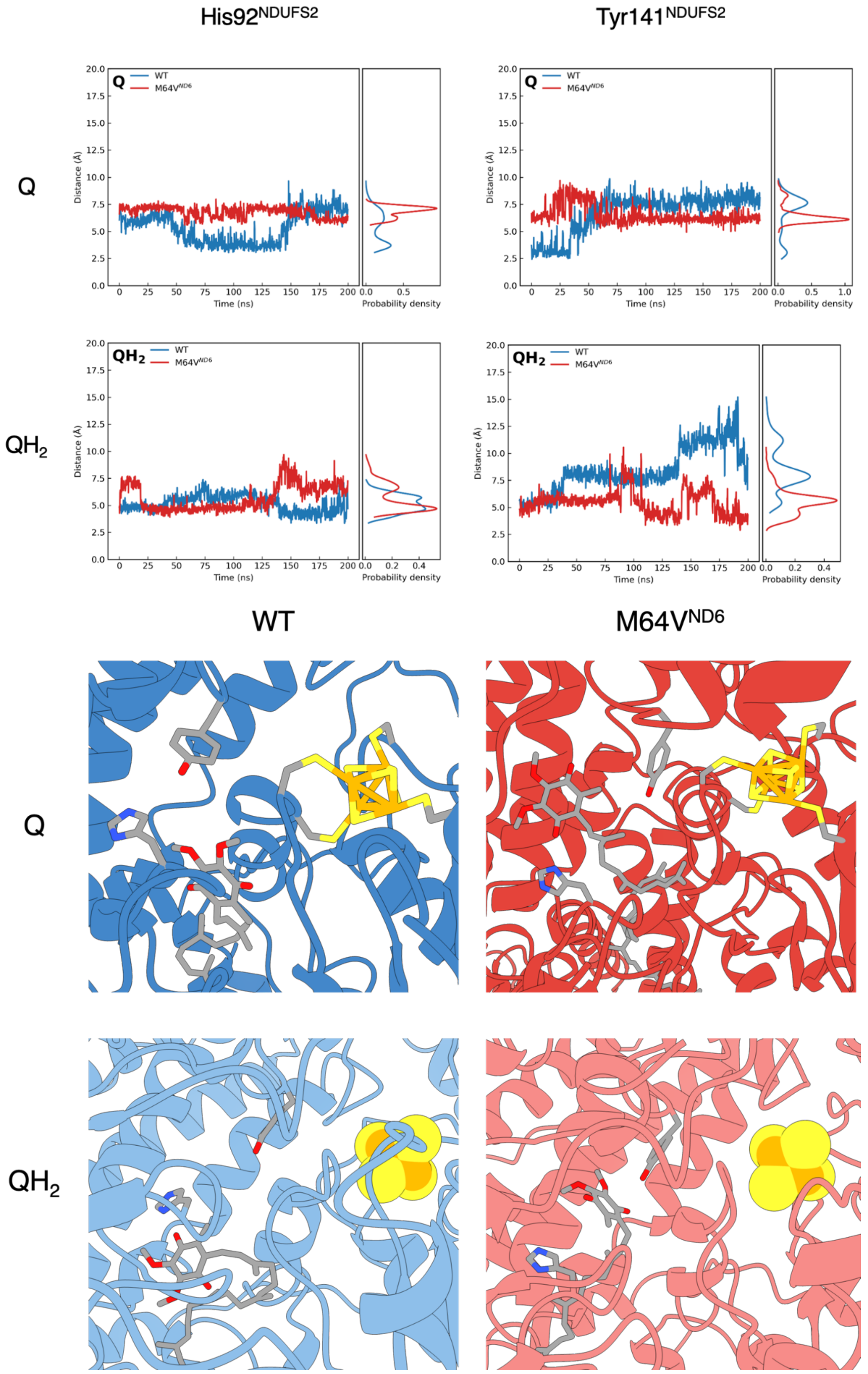


**Figure S6:** Distance between CoQ10 head group and His92$^{NDUFS2}$ (left) and Tyr141$^{NDUFS2}$ (right).

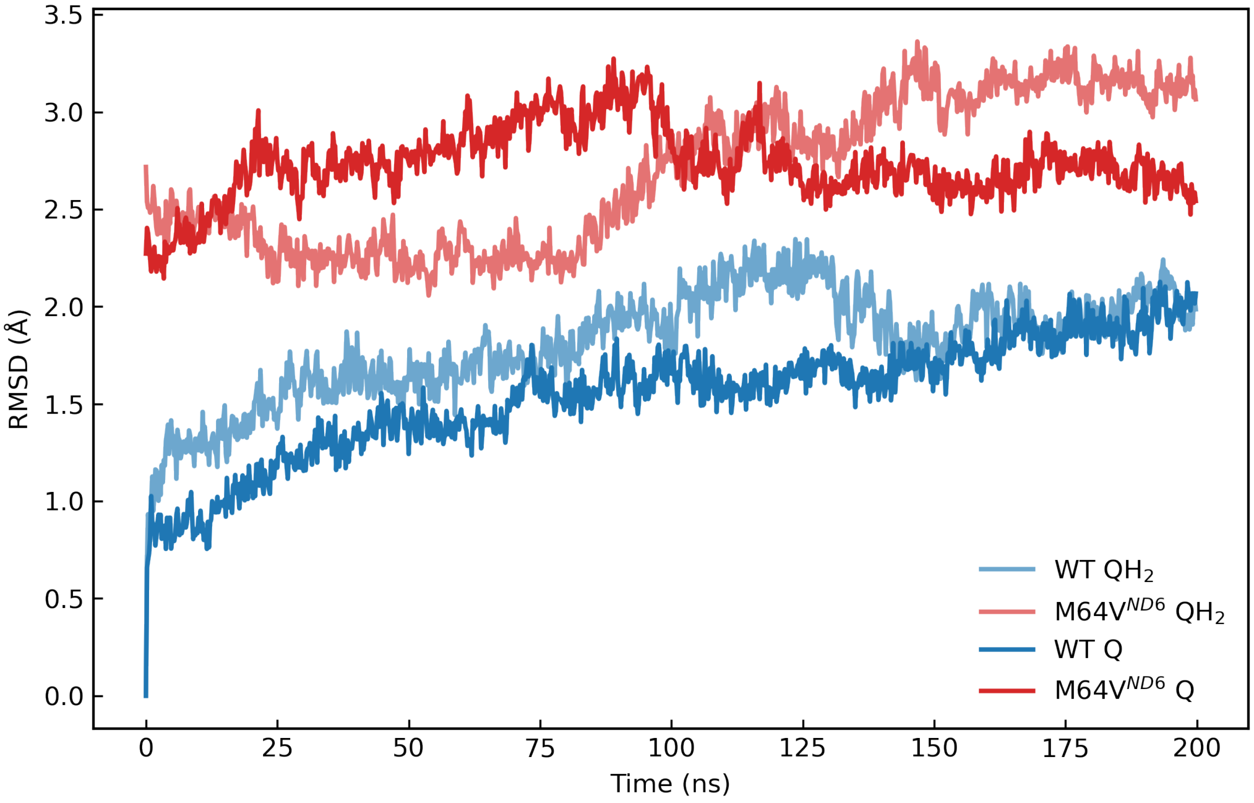


**Figure S7:** Backbone RMSD time series for ND6 in WT and ND6 mutant simulations under Q- and $QH_2$-bound conditions. For each trajectory, structures were aligned on the ND6 backbone atoms, and RMSD was calculated over the same ND6 backbone atoms. WT traces are shown in blue and mutant traces in red; $QH_2$ is indicated by a lighter tint and Q by a darker tint. Both WT and mutant RMSDs are reported relative to the WT first frame for the corresponding condition.

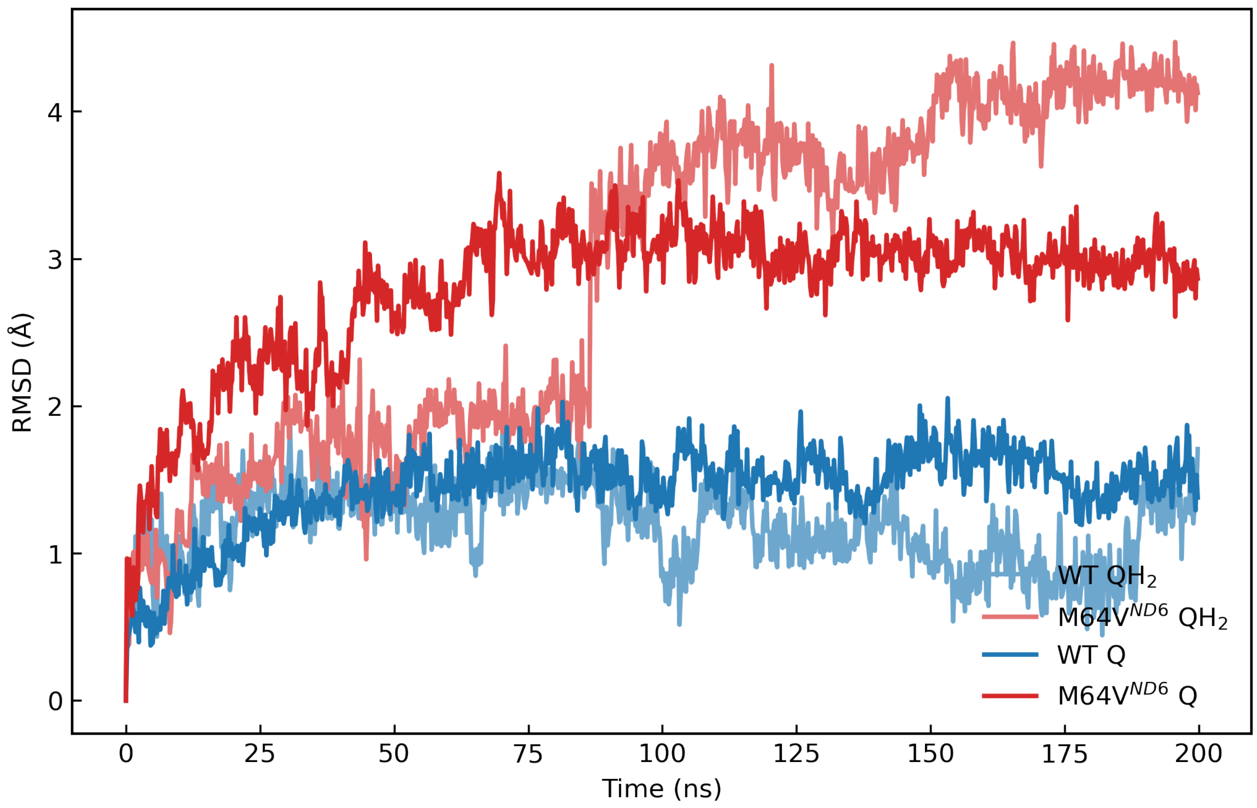


**Figure S8:** Backbone RMSD time series for the ND6 loop segment spanning residues 75–85 between TM3 and TM4 in WT and ND6 mutant simulations under Q- and $QH_2$-bound conditions. For each trajectory, structures were aligned on the backbone atoms of ND6 residues 75–85, and RMSD was calculated over the same backbone atoms. WT traces are shown in blue and mutant traces in red; $QH_2$ is indicated by a lighter tint and Q by a darker tint. WT RMSDs are reported relative to the WT first frame for the corresponding condition, whereas mutant RMSDs are reported relative to the mutant first frame for the corresponding condition.

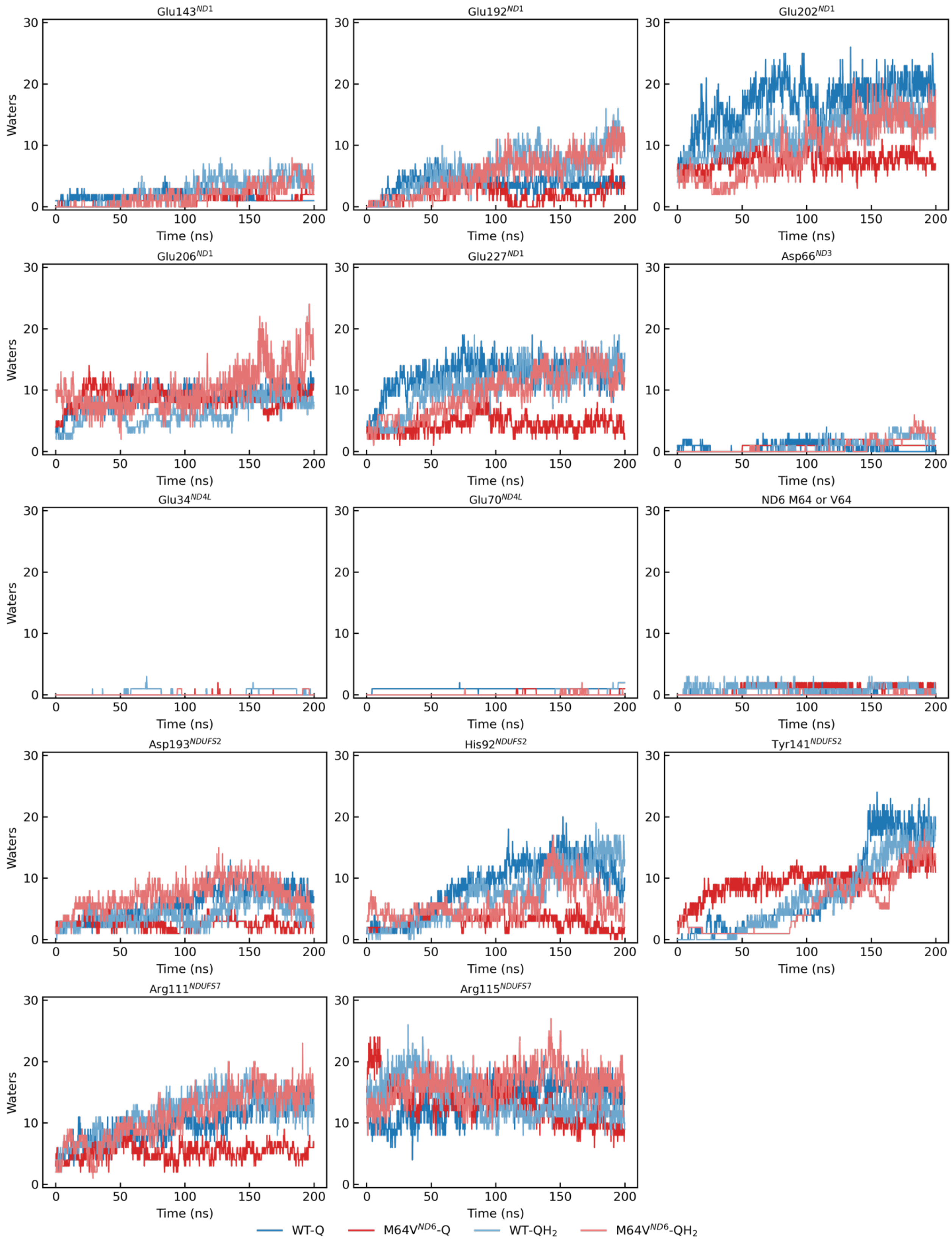
Glu143^ND1
Glu192^ND1
Glu202^ND1
Glu206^ND1
Glu227^ND1
Asp66^ND3
Glu34^ND4L
Glu70^ND4L
ND6 M64 or V64
Asp193^NDUFS2
His92^NDUFS2
Tyr141^NDUFS2
Arg111^NDUFS7
Arg115^NDUFS7
Waters
Time (ns)
WT-Q
M64V^ND6-Q
WT-QH2
M64V^ND6-QH2

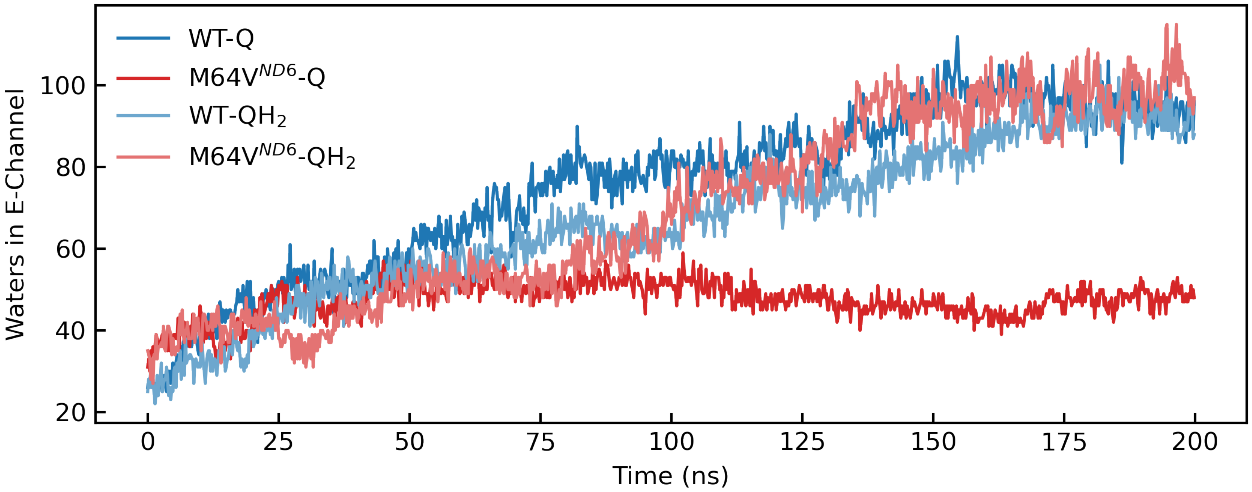


**Figure S9:** (Top) Residue breakdown of E-channel hydration for Q-bound and $QH_2$-bound equilibrium simulations. (Bottom) Total unique waters within 5 Å of said E-channel residues.

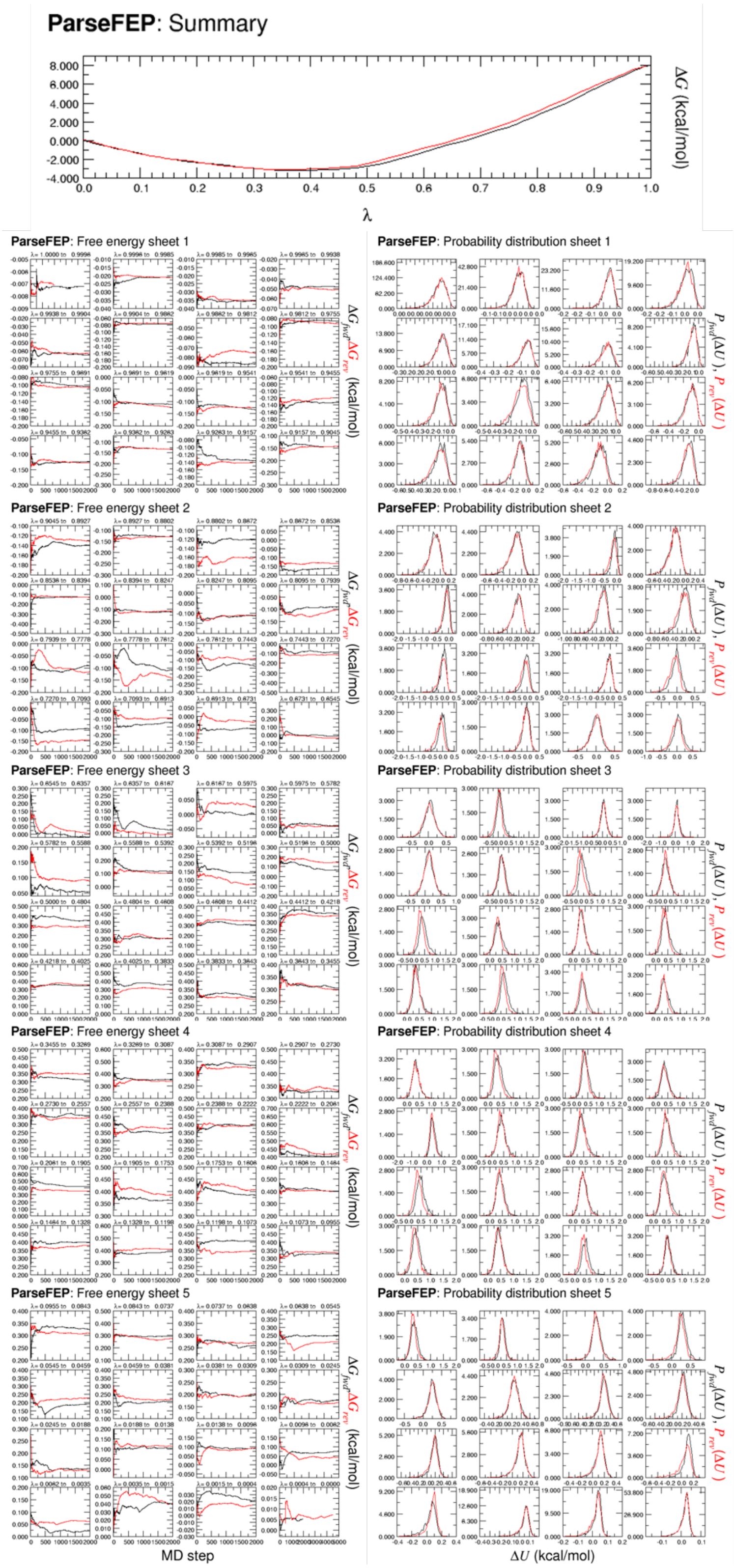


**Figure S10:** FEP Convergence using 80 windows for transformation WT-M64V in the presence of Q. N2 modeled as reduced (rN2).

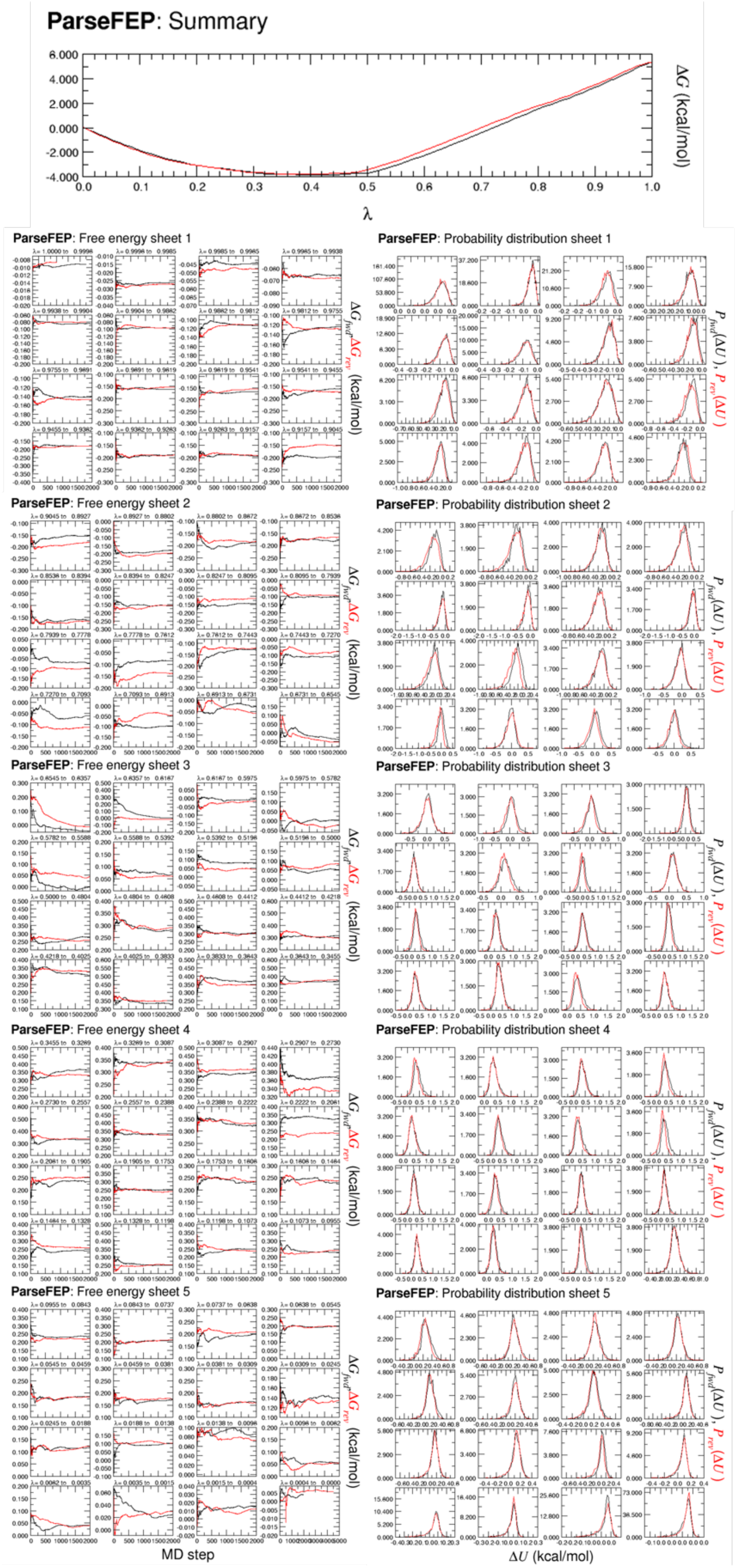


**Figure S11:** FEP Convergence using 80 windows for transformation WT-M64V in the absence of Q. N2 modeled as reduced (rN2).

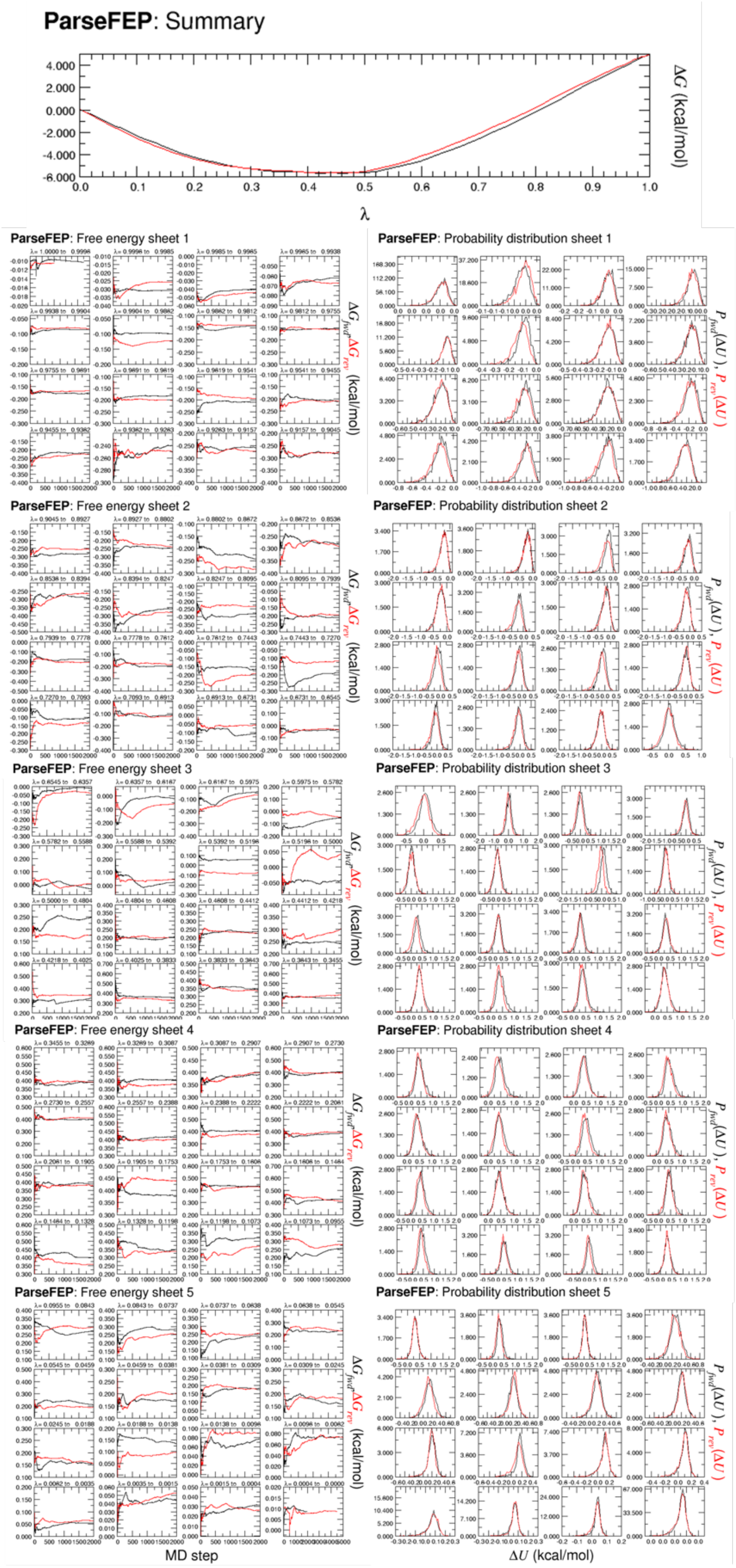


**Figure S12:** FEP Convergence using 80 windows for transformation WT-M64V in the presence of $QH_2$. N2 modeled as oxidized (oN2).

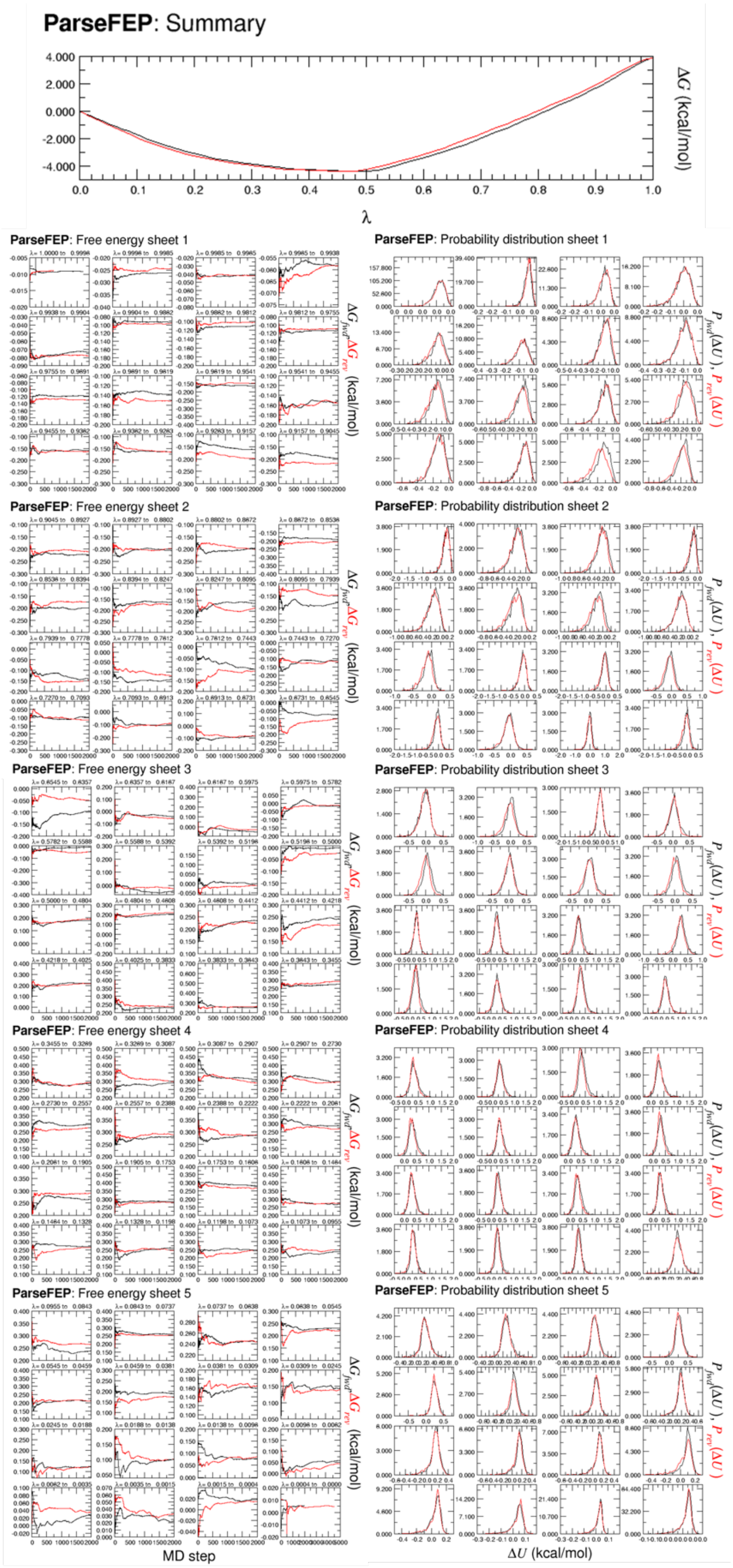


**Figure S13:** FEP Convergence using 80 windows for transformation WT-M64V in the absence of $QH_2$. N2 modeled as oxidized (oN2).
